# Delta Opioid Receptors on Parvalbumin Neurons are Necessary for the Convulsant and Anxiolytic Effects of the Delta Agonist SNC80

**DOI:** 10.1101/2025.11.14.688475

**Authors:** Marie C. Walicki, Matthew R. Mueller, Alejandrina Raigosa Garza, Erica S. Levitt, John R. Traynor, Emily M. Jutkiewicz, William T. Birdsong

## Abstract

The delta opioid receptor (DOR) is expressed broadly throughout the central and peripheral nervous systems. Activation of DOR by exogenous and endogenous ligands regulates pain, motivation, emotion and memory, but the cells and neural circuits mediating these behavioral effects remain poorly characterized. Parvalbumin-expressing (PV) interneurons express high levels of DOR transcript (*OPRD1*) and protein (Birdsong et al 2019). Parvalbumin (PV) interneurons also play a role in pain and emotional processing, suggesting that DOR signaling on PV interneurons may modulate these behaviors. To address this question we used a conditional knockout mouse line (Floxed DOR; PV-cre) to delete DOR from PV-expressing cells. First, we validated the functional loss of DOR through whole-cell electrophysiology experiments. Next, we characterized baseline behavioral phenotypes and the convulsant, pro-locomotive and anxiolytic-like behavioral responses induced by the DOR agonist SNC80 in Floxed DOR; PV-Cre mice and their littermate controls. Interestingly, we found that the convulsant and anxiolytic effects of the DOR agonist SNC80 were diminished in Floxed DOR;PV Cre animals. However, baseline behavioral phenotypes, SNC80-induced spontaneous hyperlocomotion and respiratory stimulation were conserved. These novel findings indicate that the pro-convulsant and anxiolytic effects of DOR agonists are anatomically separable from the locomotor stimulating effects and are dependent on DOR expression on PV interneurons.

## INTRODUCTION

The opioid system has been a therapeutic target for thousands of years. Opioid receptors and their endogenous opioid peptide agonists are expressed throughout the brain, where they serve as important modulators of pain, reward and emotional processing (Valentino & Volkow, 2018). While the mu opioid receptor mediates the analgesic and reinforcing effects of clinically used opioids like morphine, the delta opioid receptor (DOR) represents a potential therapeutic target for treating pain, depression and anxiety with reduced addiction liability (Pradhan et al., 2011).

Preclinical studies using both genetic and pharmacological approaches have found DOR agonists to have beneficial effects on mood, chronic pain and learning (Negus et al., 1998; Nozaki et al., 2014). However, some DOR agonists including SNC80 also produce unwanted convulsant activity (Danielsson et al., 2006). Assigning the function of DOR signaling on specific neuronal populations to specific behavioral effects is a necessary step in dissociating the desired and undesired effects of DOR agonists which is necessary for therapeutic development.

Towards this goal, global knockout of DOR results in a loss of agonist-induced hyperlocomotion, convulsions, analgesia and anxiolytic and antidepressant-like effects (Filliol et al., 2000; Gavériaux-Ruff et al., 2008; Nadal et al., 2006). Selective deletion of DOR on DLX-expressing GABAergic forebrain neurons, specifically DLX 5/6 neurons also eliminates agonist induced effects and demonstrated a novel basal anxiogenic role of DOR (Chu Sin Chung et al., 2015) and a loss of SNC80-induced seizures (Chung et al., 2015). The DLX5/6 lineage encompasses multiple types of GABA neurons including striatal spiny projection neurons and somatostatin and parvalbumin expressing interneurons (Stühmer et al., 2002). DORs are prominently expressed in all these neuron types (Chung et al., 2015; Mansour et al., 1995). However, the powerful control that PV interneurons exert over cortical and subcortical circuits makes them prime targets for behaviorally-relevant DOR action (Schwaller et al., 2004; Tahvili et al., 2025). Therefore, we aimed to determine the necessity of DOR expression on parvalbumin interneurons for the behavioral effects of the prototypical DOR agonist SNC80. To achieve this, we conditionally deleted DOR from PV neurons in mice using Floxed DOR; PV Cre mice (Gaveriaux-Ruff et al., 2011; Hippenmeyer et al., 2005; Madisen et al., 2010) and compared the basal and SNC80-induced behavioral responses with littermate control mice. We measured convulsant, anxiolytic, locomotor and respiratory activity.

## MATERIALS AND METHODS

### ANIMALS

All procedures were conducted in accordance with the National Institutes of Health guidelines and with approval from the Institutional Animal Care and Use Committee at the University of Michigan. Mice were maintained on a 12 h light/dark cycle and given ad libitum access to food and water. C57Bl/6J DORfl/fl (Gaveriaux-Ruff et al. 2011, JAX #030075) and PV-Cre mice (Hippenmeyer et al. 2005, JAX #008069) were obtained from Jackson Labs. Homozygous DORfl/fl were crossed with DORfl/fl; PV-CreWT/Cre create floxxed (Cre+) and non-floxxed (Cre-) “wild-type” littermates. Mice were 4–8 weeks at the time of stereotaxic injections, and 5-12 weeks old at the time of all electrophysiology and behavioral tests. Mice of both sexes were used. PV-cre mice were also crossed with a Cre-dependent TdTomato reporter line (AI14, Jackson Labs Stock # 007914)

### DRUGS

For behavioral studies, SNC80 (Cayman Chemical) was prepared using 0.3% HCl in sterile water for injections, the vehicle used in these experiments was 0.3% HCl in sterile water. Nitroglycerin (NTG) was prepared from a stock solution of 5.0 mg/ml NTG in 30% alcohol, 30% propelyne glycol and water (American Regent) and diluted in sterile 0.9% saline on the day of experiments. Vehicle injections for these experiments used 0.9% saline. For electrophysiology experiments, [D-Pen2,5] Enkephalin DPDPE (Tocris Bioscience) was prepared from a stock solution of 10mM in sterile water and diluted to specified concentration in 1xKREBS during slice electrophysiology experiments. Stock and working solutions of Naloxone (Tocris Bioscience) were prepared in the same manner.

### STEREOTAXIC INJECTIONS

For electrophysiology experiments, mice were injected bilaterally with an adeno-associated virus type 2 encoding channelrhodopsin [ChR2; AAV2-hsyn-ChR2(H134R)-EYFP] (UNC Vector Core, a gift from Dr. Karl Deisseroth) targeting Mediodorsal Thalamus. Mice were anesthetized with isoflurane (4% induction, 2% maintenance) and placed on a stereotaxic frame (Kopf instruments). An incision was made along the scalp and holes drilled through the skull above the injection sites. A glass pipette filled with virus was inserted into the brain and lowered to the appropriate depth. 65nl of virus was injected bilaterally into the medial thalamus (A/P, −1.2 mm; M/L, ± 0.6 mm; D/V, 3.6 mm) using a microinjector (Nanoject II, Drummond Scientific).

### ELECTROPHYSIOLOGY

Brain slices were prepared 2–3 weeks following injection of AAV. Mice were deeply anesthetized with isoflurane and decapitated. Brains were removed and mounted for slicing with a vibratome (Model 7000 smz, Campden Instruments). During slicing brains were maintained at room temperature in carbogenated Krebs’ solution containing the following (in mM): 136 NaCl, 2.5 KCl, 1.2 MgCl2–6H2O, 2.4 CaCl2–2 H2O, 1.2 NaH2PO4, 21.4 NaHCO3, 11.1 dextrose supplemented with 5 µM MK-801. Coronal sections (250–300 µM) containing the Anterior Cingulate Cortex (from the fusion of the corpus callosum to the fusion of the anterior commissure) were made and incubated in carbogenated Krebs’ solution supplemented with 10 µM MK-801 at 32°C for 30 min. Slices were then maintained at room temperature in carbogenated Krebs’ solution until used for recording.

Whole-cell recordings were made with a MultiClamp 700B amplifier (Molecular Devices) digitized at 20 kHz (National Instruments BNC-2090A). Synaptic recordings were acquired using WaveSurfer (Howard Hughes Medical Institute) and Matlab (MathWorks). Currents were evoked every 60 s by illuminating the field of view through the microscope objective (Olympus BX51WI) using a transistor-transistor logic (TTL)-controlled LED driver and a 470 nm LED (Thorlabs). The LED stimulation duration was 1 ms and power output measured after the microscope objective ranged from 0.5 to 2 mW, adjusted to obtain consistent current amplitudes across cells. The series resistance was monitored throughout the recordings, and only recordings in which the series resistance remained <15 MΩ and did not change more than 18% were included in analysis.

### CONVULSION SCORING

Mice were individually placed in plastic observation cages and monitored for 30 minutes post-injection of vehicle or drug s.c. The severity of convulsant activity was scored using a modified Racine scale (Samoriski & Applegate, 1997) adapted by (Jutkiewicz et al., 2006) and described in (Dripps et al., 2018). Convulsion severity was ranked from less severe “1” to more severe “5”. 1 includes teeth chattering, face twitching and wet dog shakes; 2-head bobbing or twitching; 3-cloni convulsion or tonic extension lasting less than 3s; 4-clonic convulsion or tonic extension lasting more than 3s; 5-clonic convulsion or tonic extension lasting more than 3s with a loss of balance. Post-convulsion catalepsy was identified by placing a rod underneath the forepaws of the animal, and a positive catalepsy score was determined by a failure to remove the forepaws within 30s and by a loss of the righting reflex. Animals that received pentylenetetrazole were sacrificed following the convulsion scoring assay due to sustained convulsions.

### ELEVATED PLUS MAZE

The elevated plus maze (EPM) was used to test anxiety-like behavior. The EPM apparatus was positioned 53 cm above the floor and consisted of 4 plexiglass arms (30 x 5 cm) extending from a center zone (5 x 5 cm). Two arms were enclosed with an opaque plexiglass wall (13 cm) and 2 other arms were open with no extending walls. Mice were individually placed in the center zone facing an open arm at the start of the 10 minute trial. The animals location, speed and entries into all zones (open arms, closed arms, and center zone) were recorded using a Firefly MV USB 2.0 camera (Teledyne FLIR integrated imaging solutions) positioned directly above the EPM and analyzed using CineLyzer behavioral tracking and analysis software (Plexon Inc).

### OPEN FIELD TESTING

The open field test (OFT) was used to test locomotion. 30 minutes prior to testing animals received vehicle or SNC80 injection (s.c). Animals were individually placed in a square plexiglass apparatus (45 x 45 cm) with 20 cm transparent plexiglass walls and under direct illumination (50 lux). Trials ran for 20 minutes and animals locomotion, speed, direction and entries into the center arena (22.5 x 22.5 cm) were recorded with Firefly MV USB 2.0 camera (Teledyne FLIR integrated imaging solutions) and analyzed using CineLyzer behavioral tracking and analysis software (Plexon Inc).

### SENSORY TESTING

The up-down method (Chaplan et al., 1994) was used to determine mechanical sensitivity with von frey filaments (0.008g to 2g). Two days prior to testing, animals were acclimated for 20 minutes in Von Frey chambers containing 5oz paper cups. On the test day animals were removed from the Von Frey chambers via the paper cups and the orbito-facial area was stimulated with Von Frey filaments to determine a baseline mechanical threshold. The 0.4g filament was used first and in the absence of a response a heavier filament (up) was tried, and in the presence of a response (face shaking, grooming, or flinching) a lighter filament was tested. Immediately after the baseline test animals were injected with 10 mg/kg nitroglycerin i.p to establish hyperalgesia. 1 hour and 15 minutes after NTG injection, animals were injected with vehicle or 10 mg/kg SNC80 s.c. Animals were placed back into the Von Frey chambers and tested for mechanical sensitivity 45 minutes after vehicle or SNC80 injection (120 minutes after NTG). All test responses after NTG were normalized to individual baseline mechanical thresholds.

### WHOLE BODY PLETHYSMOGRAPHY

Whole body respiratory measures were taken using vivoFlow whole-body plethysmography (WBP) chambers (SCIREQ Montreal, QC Canada). These chambers were supplied with 0.5L/min of medical grade air and chamber humidity, chamber temperature and animal body temperature and weights were used to calibrate volume calculations. On the day before testing animals were acclimated in the WBP chambers for 30 minutes. On test day, awake and free-moving animals were placed in chambers and data was collected and binned in 30 second intervals by Emka Technologies iox software (Sterling, VA). The duration of the test included: 5 minutes of habituation (not included in analysis), 10 minute baseline, and, after animals were injected with vehicle or 5 mg/kg SNC80 s.c., 30 minutes of post drug recording. Data analysis included frequency of breathing (Freq), tidal volume (TV), minute ventilation (MV), inspiratory time (Ti), expiratory time (Te), Ti/Te, peak inspiratory flow (PIF), peak expiratory flow (PEF), and PIF/PEF.

### IMMUNOHISTOCHEMISTRY

Parvalbumin-Cre animals crossed with an Ai14 reporter line were anesthetized with isoflurane and perfused with a 4% paraformaldehyde solution following a phosphate buffer wash. The perfused brains were kept for 12 hours at 4° until they were switched to a 1xPBS buffer for slicing. The brains were mounted for slicing 50 micron sections with a vibratome (Model 7000 smz, Campden Instruments). Brain slices were washed with a PBS buffer and counterstained with a Parvalbumin antibody (Swant). Images of the stained brain slices were taken with a CQ1 high-resolution benchtop imager (CellVoyager, Yokogawa). Images were analyzed using QuPath open-source software to identify DAPI, Parvalbumin and Ai14 positive cells within the anterior cingulate cortex AP+1.2 to AP+0.4 and dual classification parameters were set to determine the colocalization of positive detections.

### DATA ANALYSIS

One-way or two-way analysis of variance (ANOVA) with or without repeated measures (RM) were used to measure the effect of genotype and/or SNC80. All data were reported as measures ± SEM.

## RESULTS

### LOSS OF DOR IN PV NEURONS DIMINISHED THE INHIBITORY EFFECTS OF DOR SIGNALING IN THE ANTERIOR CINGULATE CORTEX

The Anterior Cingulate Cortex (ACC) receives glutamate input from the mediodorsal thalamus (MThal) and PV neurons have been shown to be exclusive mediators of feed-forward inhibitory signaling following stimulation of MThal inputs to the ACC (Delevich et al., 2015). We have previously shown that ACC PV neurons express DOR and that the DOR agonist DPDPE reduces feed-forward inhibitory synaptic transmission onto Layer V Pyramidal cells (Birdsong et al., 2019). Therefore, we measured the DOR-mediated reduction of feed-forward inhibitory signaling to confirm that Cre-expression resulted in loss of DOR function in ACC PV neurons. First, we established that the Parvalbumin-Cre mouse line effectively targets Parvalbumin positive cells through immunohistochemistry by staining ACC sections of PV-Cre;AI14 mice with anti PV antibody (Fig 1A,B). Next, we injected AAV2-ChR2 in the MThal of PV-DOR KO and WT littermate mice and we collected brain slices containing the ACC that were innervated with the thalamic ChR2-expressing terminals (Fig 1C). Whole cell voltage clamp recordings were made from Layer V ACC pyramidal neurons and MThal inputs to the ACC were optically stimulated (Fig 1D). Optically-evoked excitatory (oEPSC) and feed-forward inhibitory postsynaptic currents (oIPSCs) were recorded under baseline conditions (Fig 1E). We did not detect any significant difference in evoked EPSC or IPSC amplitudes and Excitation-Inhibition ratio (Figure 1F) between PV-DOR KO and WT littermate mice. Next, we compared the effects of the DOR agonist DPDPE (300nM) on inhibition of the oIPSC in PV-DOR KO and WT littermate mice. (Fig 1G). DPDPE significantly decreased oIPSC amplitude by an average of 49.63% ± 2.92% in WT littermates but not in PV-DOR KO (10.55% ± 2.467%) (Fig 1H). The effect of DPDPE in PV-DOR-WT was pharmacologically blocked by the DOR antagonist TIPPpsi (1 µM) indicating this is a DOR-dependent effect. Further, this reduction in inhibition by DPDPE in the presence of TIPPpsi was not statistically different from the effect of DPDPE in the PV-DOR-KO. These results demonstrate there is a functional loss of DOR-mediated inhibition of GABA release from PV interneurons in the MDThal-ACC circuit in PV-DOR-KO mice.

**Figure 1:**
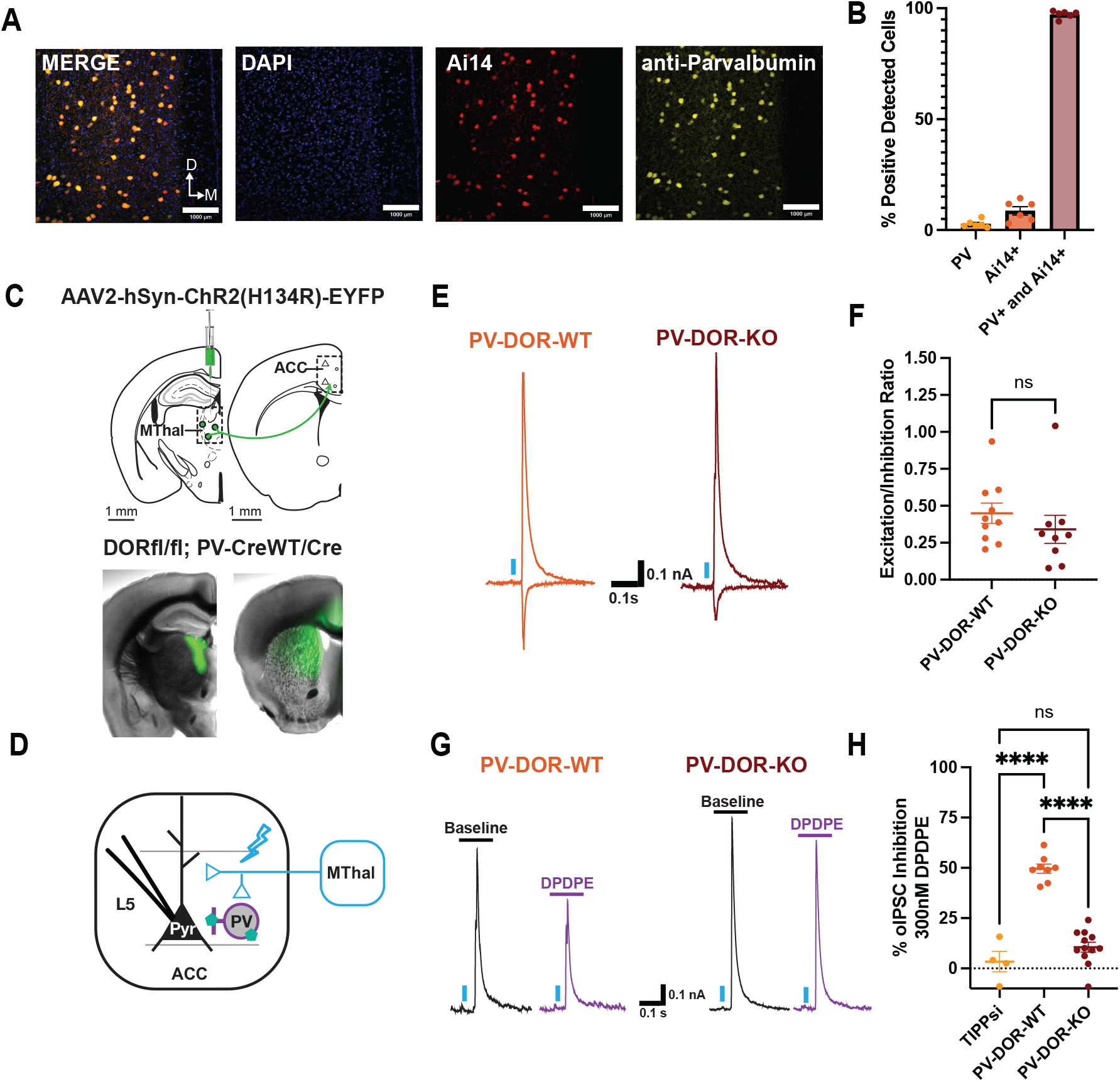
Electrophysiological characterization of floxed DOR;PV-Cre mice. (A) Representative 20x images from the ACC showing Merged overlay, DAPI (blue), Ai14 (red) and anti-Parvalbumin (yellow) from Parvalbumin-Cre;Ai14 animals. (B) Quantitative cell counting of Ai14, PV and colocalization of brain slices containing the ACC. (C) Schematic of experimental injection of channelrhodopsin injected into the mediodorsal thalamus of PV-DOR-KO and PV-DOR-WT animals and ChR2 expression in slices containing the MDThal and ACC. (D) Recording paradigm for optically evoked EPSC and IPSCs from Layer V pyramidal cells in the ACC. (E) Example traces of oEPSC and oIPSC under baseline conditions (F) Summary data of all recordings and calculated excitation-inhibition ratio under baseline conditions (Unpaired two-tailed t-test p=0.3641) (G) Example traces of oIPSCs elicited from 1ms of 470nm optical stimulation of MThal terminals in the ACC during baseline (black) and bath application of 300nM DPDPE (purple). (F) Summary data of oIPSCs from all recordings plotted as %IPSC inhibition to reflect the effect of drug application relative to baseline conditions: baseline versus 1uM TIPPsi (DOR antagonist) and 300nM DPDPE (Ordinary One-Way ANOVA p<0.0001).

### THE CONVULSANT EFFECTS OF SNC80 WERE REDUCED IN PV-DOR-KO MICE

Administration of SNC80 elicits convulsions in both rodents and primates (Danielsson et al., 2006; Hong et al., 1998). Selective deletion of DOR from GABAergic forebrain neurons reduces the convulsant action of SNC80 (Chung et al., 2015). Parvalbumin interneurons make up part of this GABAergic forebrain neuron population and their function has been implicated in seizure induction (Schwaller et al., 2004). Because PV-DOR deletion eliminated DPDPE mediated inhibition of MThal-ACC feed-forward inhibitory signaling (Fig 1E), we asked whether loss of PV-DOR could attenuate SNC80-mediated convulsions. To test this, we measured convulsant activity elicited by SNC80 at doses of 10, 32, and 100 mg/kg in PV-DOR-KO and WT littermates (Figure 2A). Using a modified Racine scale (Jutkiewicz 2006) of convulsion severity, we found a significant loss of convulsant activity at 32 and 100 mg/kg doses of SNC80 in PV-DOR-KO mice as compared to the WT littermates (Fig 2A). Additionally, 100 mg/kg SNC80 elicited a positive catalepsy score in all WT littermate mice while no PV-DOR-KO animals displayed cataleptic responses (Fig 2B). Catalepsy at the 100 mg/kg dose may indicate a shift in seizure architecture towards absence-like episodes, and this cataleptic phenotype was not observed in PV-DOR-KO mice. To determine if the loss of convulsions was specific to DOR agonists, we tested PV-DOR-KO and WT littermates with a non-opioid convulsant Pentylenetetrazol (56 mg/kg), a GABAA receptor antagonist. As expected, both PV-DOR-KO and WT littermates scored an average of 5 on the Racine scale (Fig 2C). This finding suggests that DOR expressed on PV neurons is necessary for SNC80-mediated convulsant and cataleptic activity, but not all convulsions mediated by GABA inhibition.

**Figure 2:**
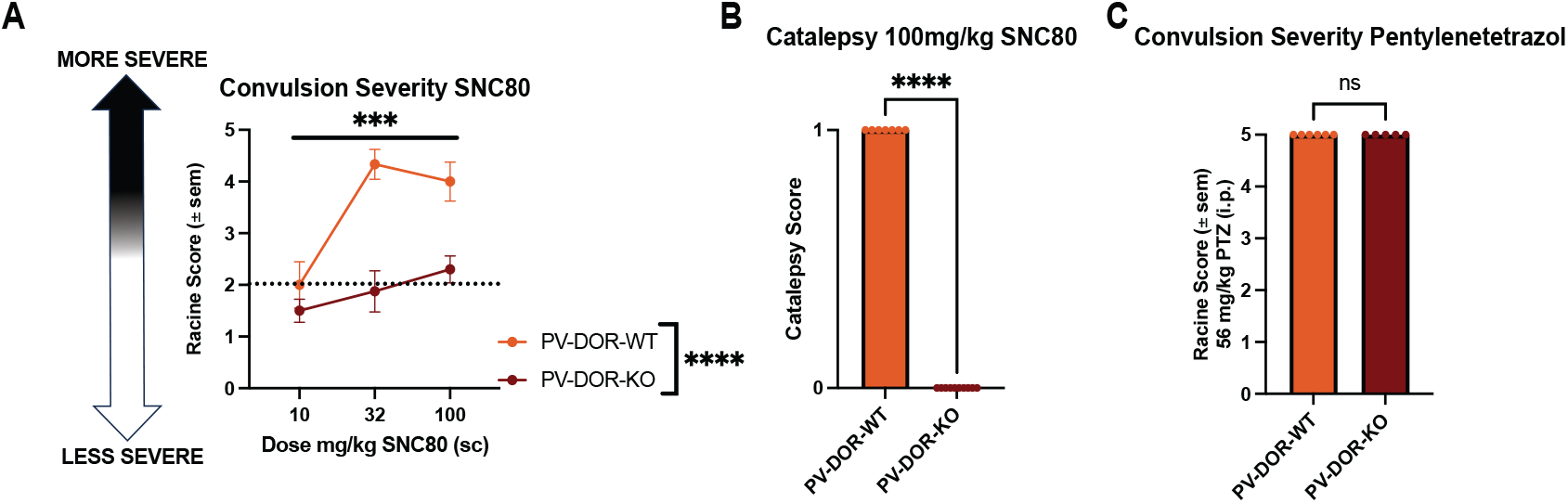
Loss of SNC80 effects of convulsion severity and type in PV-DOR-KO mice. (A) Effect of different doses of SNC80 on convulsion severity in PV-DOR-KO and PV-DOR-WT mice 10mg/kg n=6,6, 32 mg/kg n=and 100mg/kg (Two-Way ANOVA drug P=0.0004, genotype P<0.0001, interaction P=0.0281) (B) Catalepsy scores of PV-DOR-WT and PV-DOR-KO mice from 100mg/kg SNC80 (Mann-Whitney test P<0.0001) (C) Effect of Pentylenetetrazol (56 mg/kg) on convulsion severity in PV-DOR-WT and PV-DOR-KO mice (Mann-Whitney test P>0.9999)

### THE ANXIOLYTIC EFFECTS OF SNC80 WERE DIMINISHED IN PV-DOR-KO MICE

SNC80 has well-established anxiolytic effects (Perrine et al., 2006) while a loss of DOR in GABAergic forebrain neurons has a paradoxical anxiogenic role (Chu Sin Chung et al., 2015; Filliol et al., 2000). Therefore, we used the elevated plus maze to measure basal anxiety-like behavior and SNC80 induced anxiolytic effects across genotypes. SNC80 increased the time spent in the open arms in WT littermates suggestive of an anxiolytic effect of SNC80 (Fig 3B). However, this increase in time in open arms was reduced in PV-DOR-KO mice. While there was a main effect of genotype on both the time and entries into the open arms, the average speed and track length increased with SNC80 across genotypes (Fig 3 D,E). This suggests that the locomotor stimulating effects of SNC80 were conserved in PV-DOR-KO.

**Figure 3:**
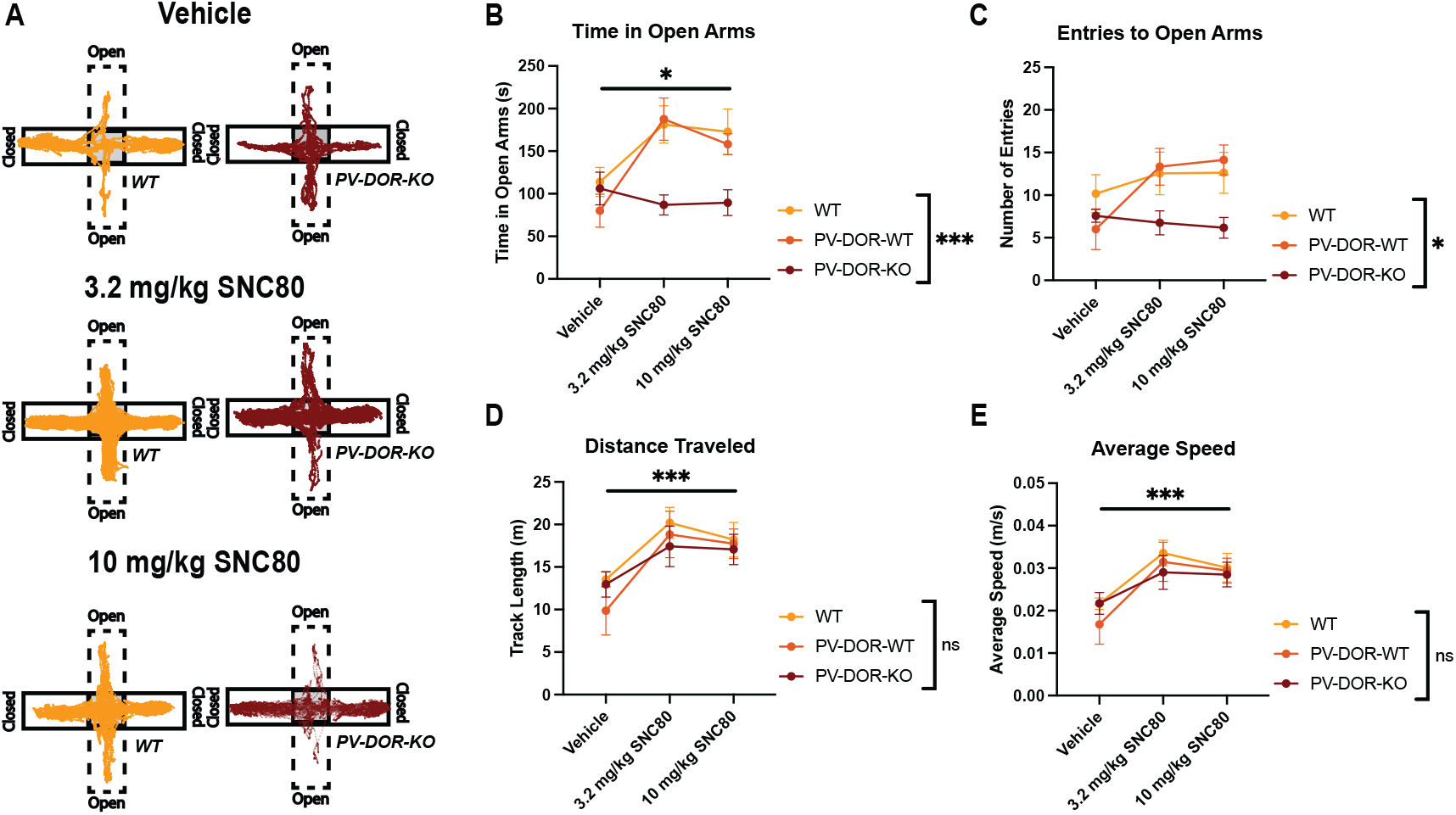
Loss of SNC80 effects on anxiety-like behaviors measured through elevated plus maze. (A) Representative traces from elevated plus maze of WT (n=28,(PV-DOR-WT (n=18), and PV-DOR-KO (n=21) animals different doses of SNC80. (B) raw time in center zone (Two-Way ANOVA drug P=0.0117, genotype P=0.0010, interaction P=0.0567) (C) entries to center (Two-Way ANOVA drug P=0.1697, genotype P=0.0105, interaction P=0.3458) (D) distance traveled (Two-Way ANOVA drug P=0.0003, genotype P=0.446, interaction P=0.84) (E) average speed (Two-Way ANOVA drug P=0.0002, genotype P=0.5712, interaction P=0.8417) from WT, PV-DOR-WT and PV-DOR-KO mice under vehicle, 3.2mg/kg and 10mg/kg SNC80. Two-Way ANOVA drug, genotype and interaction significance are denoted * P<0.05, **P<0.001, ***P<0.0001, and ****P<0.00001.

### THE LOCOMOTOR STIMULATING EFFECTS OF SNC80 WERE MAINTAINED IN PV-DOR-KO MICE

As seen in the elevated plus maze, SNC80 and other DOR agonists stimulate locomotor activity in rodents (Jutkiewicz et al., 2008). Prior work has demonstrated that DOR action in the striatum or ventral tegmental area may be responsible for the locomotor-stimulating effects of DOR agonists (Bosse et al., 2008). However, the specific sites of DOR action which contribute to the enhanced locomotor effects of DOR agonists remain to be identified. We previously observed an increase in track length and speed across genotypes in EPM (Fig 3). To further investigate the effect of PV-DOR-KO on motor stimulation we used Open Field Testing (OFT) to measure baseline locomotion and anxiety-like phenotypes in PV-DOR-KO mice compared to WT littermates, and C57Bl/6J “WT” mice (Fig 4). We found no measurable difference in track length, time in center, entries to center, or speed between groups at baseline. Next, we measured the effects of SNC80 in OFT measures (Fig 4B-E). Animals received a vehicle, 3.2 or 10 mg/kg s.c dose of SNC80 or vehicle 30 minutes prior to OFT. Two-way ANOVA [F(2,49)=0.697] revealed no significant main effects of genotype on any reported measure, but a significant main effect of SNC80 [F(2,49)=53.17] in distance traveled and average speed. This suggests that the pro-locomotive effects of SNC80 are maintained in PV-DOR-KO mice. Time in center, a traditional measure of anxiety-like behavior, did not increase in any group relative to vehicle-treated controls (P>0.05), but rather decreased in all groups with SNC80 on board. The significant increase in speed and track length with SNC80 treatment demonstrates that OFT may be less robust for testing the anxiety-relieving components of DOR agonists when there is such a large increase in locomotor activity.

**Figure 4:**
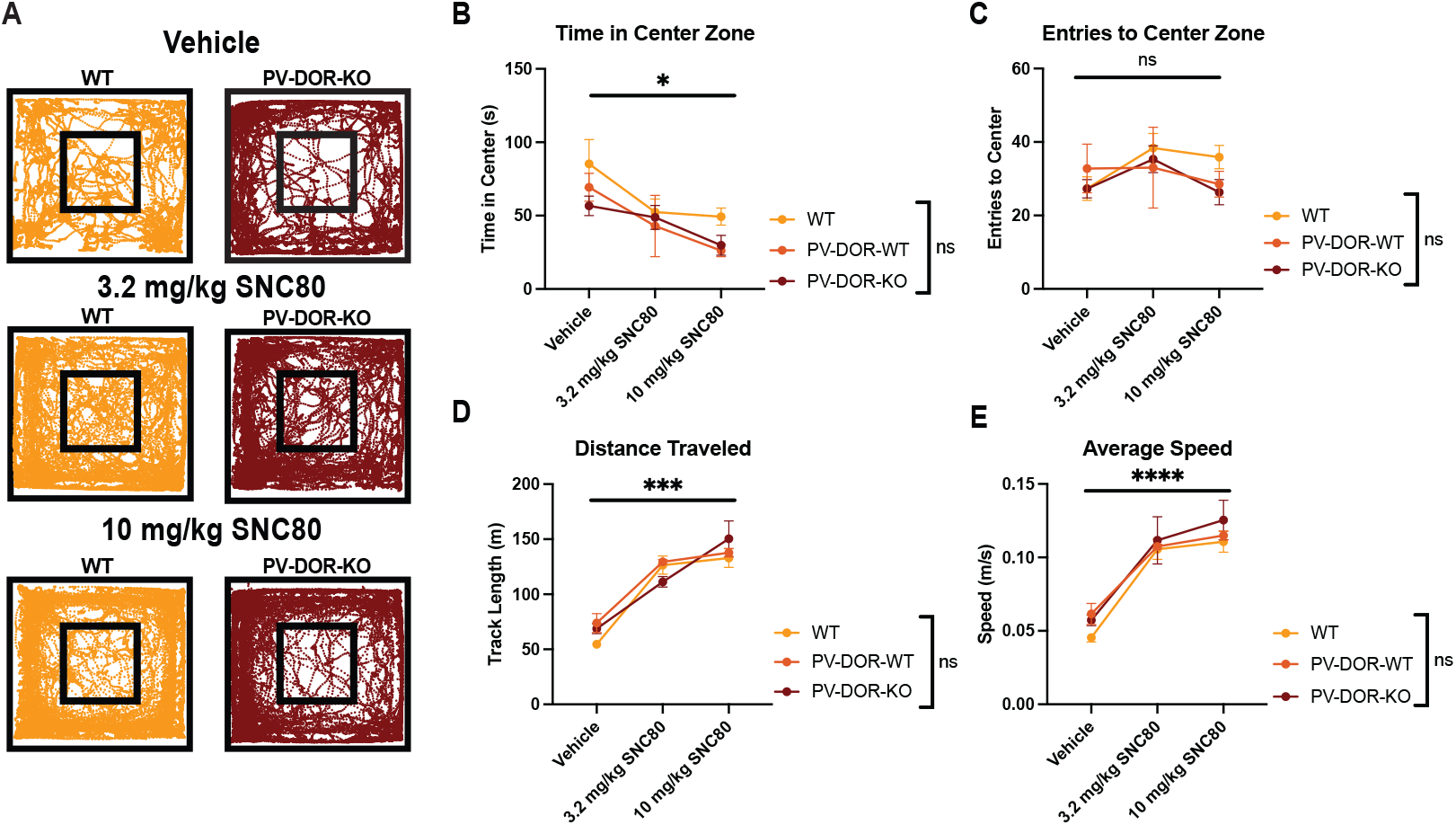
SNC80 effects on locomotor stimulation measured with open field testing. (A) Representative tracks of WT (n=28), PV-DOR-WT (n=11) and PV-DOR-KO (n=20) animals in OFT with different doses of SNC80 (B) raw time in center zone (C)entries to center zone (D) distance traveled and (E) average speed. Two-Way ANOVA drug, genotype and interaction significance are denoted * P<0.05, **P<0.001, ***P<0.0001, and ****P<0.00001.

### SNC80 INCREASED RATE OF BREATHING IN PV-DOR-KO MICE

DOR agonists have been shown to stimulate respiration as measured by an increase in breathing rate (Hug et al., 2022) but the site of DOR action in respiratory circuits is unknown. Because we saw enhanced SNC80-induced locomotor stimulation in elevated plus maze (Fig 3) and open field testing (Fig 4), we tested whether the PV-DOR-KO mice have alterations in frequency of breathing under normal conditions and following SNC80 (5mg/kg). We observed no significant changes in breathing frequency at baseline conditions across genotypes (Fig 5A). Additionally, SNC80 injection increased frequency of breathing to a similar extent across all genotypes compared to vehicles (Fig 5B). These results indicate that, similar to locomotor stimulation, PV-DOR is not necessary for SNC80-induced respiratory stimulation.

**Figure 5:**
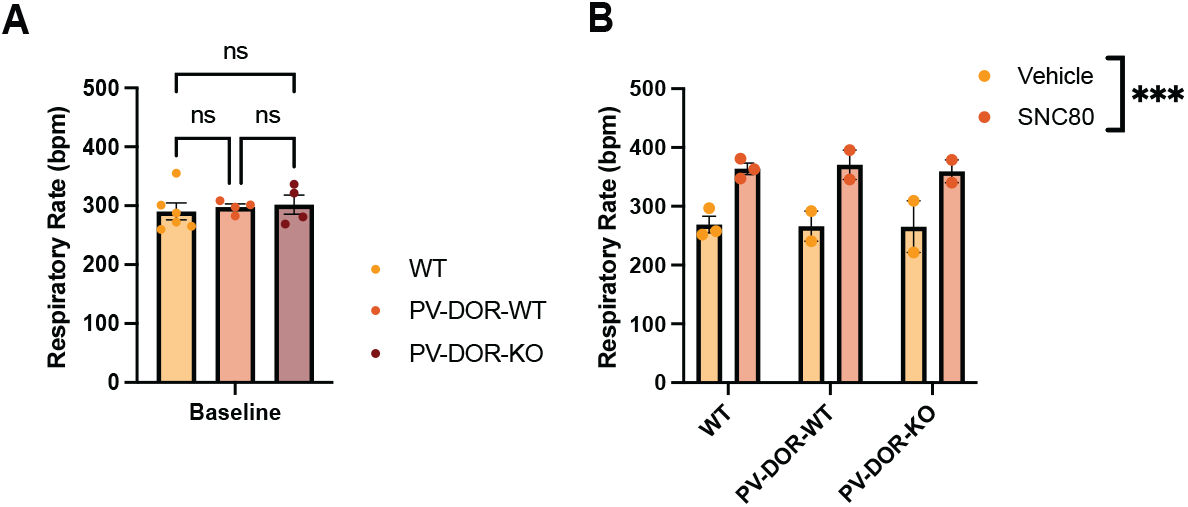
SNC80 effects on stimulation of breathing. (A) Respitrator rate (breaths per minute, bpm) measured under baseline conditions (One-Way ANOVA P=0.8236) (B) frequency of breathing following vehicle or SNC80 conditions in WT, PV-DOR-WT and PV-DOR-KO animals (Two-Way ANOVA drug P=0.007, genotype P=0.9672, interaction P=0.9691)

### SENSORY SENSITIVITY WAS MAINTAINED IN PV-DOR-KO MICE

Parvalbumin neuron activity and DOR agonists have been shown to mediate pain-related behaviors such as nociception and the development of hyperalgesia (Cahill et al., 2022; Nadal et al., 2006; Qiu et al., 2024). DOR agonists have been shown to be effective in attenuating nitroglycerin-induced hyperalgesia, a model of migraine-like pain. This effect was lost when DOR was knocked out of DLX-expressing neurons (Pradhan et al., 2014). We used the nitroglycerin-induced hyperalgesia assay combined with orbito-facial von Frey testing to determine whether PV-DOR-KO affected the development of hyperalgesia. Additionally, we tested whether SNC80 had anti-hyperalgesic effects in PV-DOR-KO mice. There were no statistically significant differences in baseline mechanical sensitivity (Fig 6A) across genotypes. Intraperitoneal injections of 10mg/kg nitroglycerin significantly decreased the cephalic mechanical threshold in vehicle injected animals relative to individual baseline measurements in both WT and PV-DOR-KO mice. Additionally, this NTG-induced decrease in mechanical thresholds was prevented by SNC80 treatment in both WT and PV-DOR-KO mice (Fig 6B). These results indicate that PV-DOR-KO animals develop hypersensitivity following NTG administration at similar severity levels. Additionally, the anti-hyperalgesia effects of SNC80 were conserved across genotypes.

**Figure 6:**
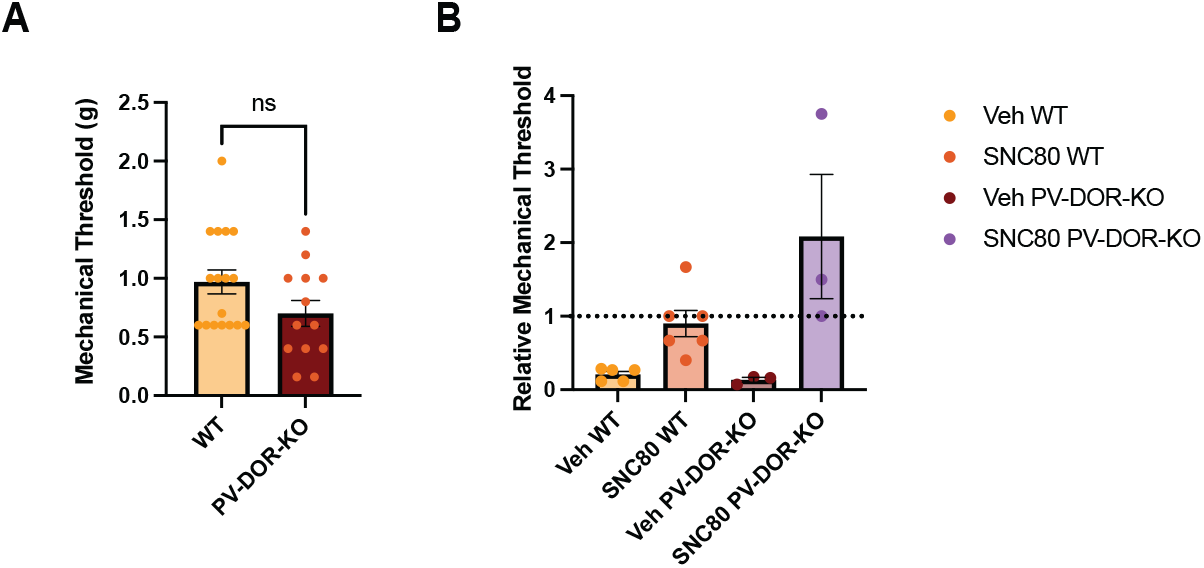
Antihyperalgesic effects of SNC80 following NTG treatment. (A) Baseline mechanical withdrawal threshold (g) in wild-type (WT) and floxedDOR;PV-Cre (PV-DOR-KO) animals (two-tailed t-test P=0.0934) (B) Effects of vehicle or SNC80 treatment on mechanical withdrawal thresholds calculated relative to individual baselines in NTG-treated wild-type and floxed DOR;PV-Cre (PV-DOR-KO) animals (Two-Way ANOVA drug P=0.0013, genotype P=0.1112, interaction P=0.0745)

## DISCUSSION

In this study, we used Floxed DOR;PV Cre mice to selectively delete delta opioid receptors from parvalbumin interneurons. Using electrophysiological approaches, we observed a loss of DOR signaling on parvalbumin cells and diminished DOR inhibition of feedforward inhibition in the ACC of PV-DOR-KO animals. By characterizing the behavioral effects of PV-DOR-KO, our study presents a novel finding that the convulsant and anxiolytic-like effects of the DOR agonist SNC80 are separable from the locomotor-stimulating and analgesic effects, with selective DOR expression on PV neurons being necessary only for convulsant and anxiolytic-like effect. DOR is widely expressed on PV neurons. Therefore, further studies are needed to identify which specific PV neurons contribute to these respective behavioral effects of DOR agonists.

We observed a loss of full myoclonic convulsions (Racine scores of 4 or 5) in PV-DOR-KO animals. While we observed a reduced Racine score in our PV-DOR-KOs, not all pre-convulsant activity was eliminated. Our observations suggest that there are other sites of DOR action that contribute to SNC80-induced pre-convulsant activity. While PV-DOR-KOs did not eliminate all SNC80-induced pre-convulsant activity, we observed no cataleptic, or “absence-like” seizure events at high doses of SNC80 (Fig 2B). A prior study knocking out DOR from forebrain GABA neurons, using Floxed DOR;Dlx-Cre mice, found a complete loss of all SNC80 induced convulsant activity measured through EEG (Chung et al., 2015). While myoclonies were not observed in Dlx-DOR animals, SNC80 increased spike-and-wave discharges, a measure of absence-like seizures. Because PV interneurons are a subpopulation of GABA forebrain neurons, our findings suggest that PV-DOR may contribute to SNC80-induced absence-like seizure activity as well as full myoclonic convulsions while other neuronal populations such as somatostatin interneurons or striatal spiny projection neurons may further affect pre-convulsant activity. Parvalbumins cells are fast-spiking GABAergic cells and are critical modulators of synaptic transmission and gamma-band oscillations (Tahvili et al., 2025). Parvalbumin cells in the cortex regulate network excitability and a loss or hypofunction of parvalbumin cells impairs local inhibition which increases network excitability. These effects increase seizure severity and susceptibility in rodents (Schwaller et al., 2004). Inhibition of PV transmission by DOR activation likely would result in similar effects. Parvalbumins across the cortex and hippocampus express high levels of DOR transcripts, so DOR inhibition of PV cells within these regions may impair local inhibitory circuitry and result in increased seizure-like activity. Further studies using EEG or intrinsic optical imaging along with selective DOR KO may further elucidate roles of DOR on GABA neuron subpopulations of the Dlx lineage.

Perhaps the most intriguing finding of our study is the loss of the anxiolytic effects of SNC80 in the elevated plus maze test in PV-DOR-KO animals. Prior studies of selective DOR knockdown on GABAergic forebrain neurons elicits a novel low-anxiety phenotype (Chu Sin Chung et al., 2015), which is opposite the observed elevated anxiety state of global DOR knockout animals. This initial finding suggests that DOR action on forebrain GABAergic neurons may be involved in regulating anxiety states basally. PV-DOR-KO animals did not exhibit changes in baseline anxiety states as measured through baseline responses in OFT and EPM. However, the anxiety-relieving effect of SNC80 was lost (Figure 3). Studies of reduced or eliminated parvalbumin function also result in increased anxiety states (Perlman et al., 2021). Specifically, PV-DOR forebrain neurons may be contributing to the emotion regulating effects of DOR signaling through both PV interneurons and other GABAergic neuronal populations. However, further studies are needed to determine which DOR-expressing interneurons modulate basal emotional states and which PV neurons contribute to the anxiolytic effects.

Along with their anxiety-relieving effects, DOR agonists have been well characterized to have antidepressant-like action in rodents (Jutkiewicz et al., 2005). DOR knockdown studies cited in this work have reported to have an enhancement of reward and motivation behaviors (Filliol et al., 2000). Our studies did not examine despair-like or motivation behaviors, but as our findings report a change in anxiety-like effects, we may also see alterations in these other emotion-related behaviors. Further characterization of these behaviors within our model may help to understand how PV-DOR signaling contributes to the beneficial aspects of DOR.

Constitutive DOR knockout mice demonstrate enhanced spontaneous locomotion, but this effect was absent in the Dlx-DOR-KO model (Chu Sin Chung et al., 2015). Similar to Dlx-DOR-KO, PV-DOR-KO mice did not display enhanced spontaneous locomotion during baseline observations in OFT. Further, SNC80-induced hyperlocomotion was similar to control mice. In fact, SNC80 significantly increased track length and speed in both OFT and EPM assays to a similar extent in PV-DOR-KO, PV-DOR-WT control and WT control animals. Previous work has suggested that DOR action on striatal spiny projection neurons play a prominent role in locomotor activity induced by DOR agonists (Bosse et al., 2008), which is consistent with our present study.

In addition to the mood altering effects, DOR agonists elicit analgesia (Cahill et al., 2022; Negus et al., 1998). In rodent models of migraine-like pain, DOR agonists effectively treat allodynia associated with migraine-like pain better than other opioid receptor agonists (Pradhan et al., 2014). We found that our PV-DOR-KO animal developed hyperalgesia induced by nitroglycerin at a similar rate and severity to wild-type mice. This finding suggests that a loss of DOR function on PVs is not associated with alterations in migraine-like pain transmission. Endogenous DOR function is thought to modulate the severity of pain as well. Constitutive DOR knockout studies have shown enhanced mechanical and thermal pain responses in DOR KO animals compared to WTs (Gavériaux-Ruff et al., 2008; Nadal et al., 2006). However, PV-DOR-KO mice did not show a significant enhancement in basal mechanical sensitivity using the orbitofacial von Frey test. Further, while SNC80 blocked the hyperalgesic effect of NTG, we did observe a relatively stronger effect of SNC80 in PV-DOR-KO as compared to WT animals. This suggests that while baseline hyperalgesia is not changed across genotypes, the effect of SNC80 on anti-hyperalgesia may be altered. Parvalbumins cells in the central and peripheral pathways of gate pain transmission through inhibition (Qiu et al., 2024). Disinhibition of PVs by DOR may alter PV transmission in a similar manner, and this disruption of disinhibition by PV-DOR-KO may result in the enhanced anti-hyperalgesic effect of SNC80 observed in our study. Further characterization of PV-DOR-KO model on the development of chronic pain, and analgesic effects of opioid agonists should be performed.

Understanding the neural targets of DOR signaling that give rise to DOR-mediated behaviors are crucial for developing clinically safe and effective DOR therapeutics for depression, anxiety and pain. This work highlights a novel selective knockdown of DOR on PV cells modulates distinct DOR-mediated behaviors. This work will allow for further investigation of these parvalbumin interneuron-dependent and independent sites of DOR action and potentially facilitate the development of DOR-targeting therapeutics.

## FOOTNOTES

This authors declare no competing interests.

This work was supported by National Institutes of Drug Abuse Grant T32DA007281 (to M.C.W.) and National Heart Lung and Blood Institute Grant R01HL174547 (to E.S.L).

